# On the mechanism for winter stem pressure build-up in walnut trees

**DOI:** 10.1101/2023.12.21.572762

**Authors:** Cyril Bozonnet, Marc Saudreau, Eric Badel, Guillaume Charrier, Thierry Améglio

**Affiliations:** Université Clermont Auvergne, INRAE, PIAF, 63000 Clermont-Ferrand, France

**Keywords:** Embolism recovery, Modelling, Sugar transport

## Abstract

Xylem embolism is a significant factor in tree mortality. Restoration of hydraulic conductivity after massive embolisation of the vascular system requires the application of positive pressure to the vessels and/or the creation of new conductive elements. Some species generate positive pressure from the root system to propagate pressure in distal, aboveground, organs in spring, whereas other species generate positive pressure locally at the stem level during winter. We provide a mechanistic explanation for winter stem pressure build-up in the walnut tree. We have developed a physical model that accounts for temperature fluctuations and phase transitions. This model is based on the exchange of water and sugars between living cells and vessels. Our computations demonstrate that vessel pressurization can be attributed to the transfer of water between vessels across the parenchyma rays, which is facilitated by a radial imbalance in sugar concentration. The ability to dispose of soluble sugars in living cells, and to transport them between living cells and up to the vessels, are identified as the main drivers of stem pressure build-up in the walnut tree.

## Introduction

Massive embolism in xylem causes a decline in trees’ hydraulic conductivity and is among the factors causing their death (Sperry and Tyree, 1988; Brodribb and Cochard, 2009; Mantova et al., 2022). This embolism can occur as a result of either drought, leading to air seeding in xylem conduits (Hargrave et al., 1994; Choat et al., 2016), or due to freeze-thaw cycles (Sperry and Sullivan, 1992; Charra-Vaskou et al., 2016). Despite previous studies (Salleo et al., 1996; Nardini et al., 2011), hydraulic conductivity seemingly cannot be re-established while the xylem remains under tension (Charrier et al., 2016). Hydraulic conductivity recovery happens through creating new conducting vessels (Cochard and Tyree, 1990), pressurizing xylem conduits to remove air bubbles (Sperry et al., 1987; Hacke and Sauter, 1996), or a combination of both (Cochard et al., 2001; Améglio et al., 2002). In species that pressurize their xylem conduits, it is important to differentiate between species where the pressure comes only from the roots in spring (Fisher et al., 1997), and species where the pressure can also come from the stem in winter (Améglio et al., 2001). The walnut tree has the ability to use both strategies to generate positive pressure (Ewers et al., 2001).

Note that for many species during winter, there are pressure variations associated with freezing events (Robson and Petty, 1987; Milburn and O’Malley, 1984; Améglio et al., 2001). They are related to phase changes and freeze-induced water fluxes (Ceseri and Stockie, 2013; Graf et al., 2015; Bozonnet et al., 2023; Zarrinderakht et al., 2024). What we call here “stem pressure” is a pressure that stays positive in a stem conduit even after sap thawing and that could subsequently lead to embolism recovery by refilling, once capillary forces between an embolized and a pressurized conduit are overcome. The term “build-up” refers to the gradual increase of this pressure with time.

Winter stem pressure has been extensively studied for walnut tree (Améglio and Cruiziat, 1992; Améglio et al., 2001; Ewers et al., 2001; Améglio et al., 2002, 2004), and maple tree (Milburn and O’Malley, 1984; Tyree, 1983; Cirelli et al., 2008; Ceseri and Stockie, 2013; Graf et al., 2015; Zarrinderakht et al., 2024). Note that the work of Graf et al. (2015) also includes a comparison with experimental results for walnut tree. In this work, we focus on walnut tree.

Walnut winter stem pressure has been studied using laboratory experiments in (Améglio et al., 2001) that demonstrated many features associated with this phenomenom: 1) the pressure can be generated for stems disconnected from the rest of the tree; 2) the pressure rise starts at positive temperature and stops at higher temperature (typically *>* 5°C); 3) the pressure rises much more during successive freeze-thaw cycles rather than during continuous exposure to low and non-freezing temperature; 4) at the end of the experiments, the magnitude of the pressure build-up is positively correlated with the osmolarity of the xylem sap; 5) earlier stem defoliation and exposure to high temperature (typically 18°C) before successive freeze-thaw cycles both reduce the xylem sap osmolarity and xylem pressure.

During winter, when sap mineral content is low, xylem sap osmolarity depends on a balance of sugar fluxes between vessels and vessel-associated cells (VACs, Améglio et al. (2004)), mediated by temperature dependent H^+^/sugar co-transport (Alves et al., 2007). Particularly, sugar fluxes from VACs to vessels are thought to be diffusive (facilitated by a specific protein), whereas a H^+^/sugar co-transport, related to the ATP-ase activity, drives the fluxes from vessels to VACs at sufficiently high temperature (Améglio et al., 2004; Decourteix et al., 2008). Pressure changes in the thawed state are thus believed to be due to the exchange of water between VACs and vessels, which are triggered by the corresponding sugar fluxes.

In this work, we built a mechanistic model for pressure build-up in walnut tree that is intended to reproduce the specific features listed above. We particularly explored the link between xylem pressure rise and xylem sap osmolarity.

To do so we have developed a comprehensive physical model that integrates water and heat fluxes, phase changes, pressure-volume relationships, and sugar fluxes within various tissues. The model incorporates changes in stem diameter that are related to water flows between living cells (in bark or xylem tissues) and apoplast. During freeze-thaw cycles, pressure and stem diameter changes are inter-related as we demonstrated in our previous work (Bozonnet et al., 2023). This feature allows future comparison of the model results with non-invasive measurements of stem diameter changes.

After a presentation of the effects of sugar fluxes on pressure changes, we explored the role of sugar permeabilities and initial concentration on pressure build-up. We then compared the outputs of the model (pressure level, xylem sap osmolarity) to the experimental results of Améglio et al. (2001). We finally presented a thorough analysis of the model results, highlighting key findings and potential avenues for further research. The model is freely available along with the paper so that other scientists could benefit from its use and contribute to its development.

## Material and methods

### General description of the numerical model

The present model is a modified version of a previous one presented and validated in Bozonnet et al. (2023). This model was based on earlier modeling efforts about pressure changes in maple trees (Ceseri and Stockie, 2013; Graf et al., 2015). We provide here a general description of the model and highlight its difference with its previous version.

The model relies on a wood anatomy description in the transversal plane of a wood section, as described by Alves (Alves et al., 2007). The structure of our model (figure 1) groups the essential anatomical elements to simulate the processes described above: xylem vessels, vessel-associated cells (VACs) and bark cells. It shares some similarities with the model of Hölttä et al. (2006): living cells are interconnected by a parenchyma ray, connected at its periphery to the bark cells, and connected to a radial alignment of vessels. We assume that the external temperature field is homogeneous around the stem, i.e. axi-symmetric, so that radial exchanges are only modelled along one ray and rescaled at the xylem/bark interface by *N*_*ray*_, the number of parenchyma rays. The longitudinal dimension is not considered. The parenchyma ray itself is not described explicitly, i.e., individual (isolated) ray cells are omitted, but rather represented by a hydraulic resistance between VACs, and between VACs and bark cells. Water flows and volume changes are computed for one VAC per vessel and for one bark cell in the bark tissue. These water flows and volume changes are then rescaled by *N*_*vac*_, the number of VACs connected to each vessel, and *N*_*bark cell*_, the number of living cells in the bark, similarly to what is done in Graf et al. (2015) for the fiber/vessel fluxes.

**Figure 1:**
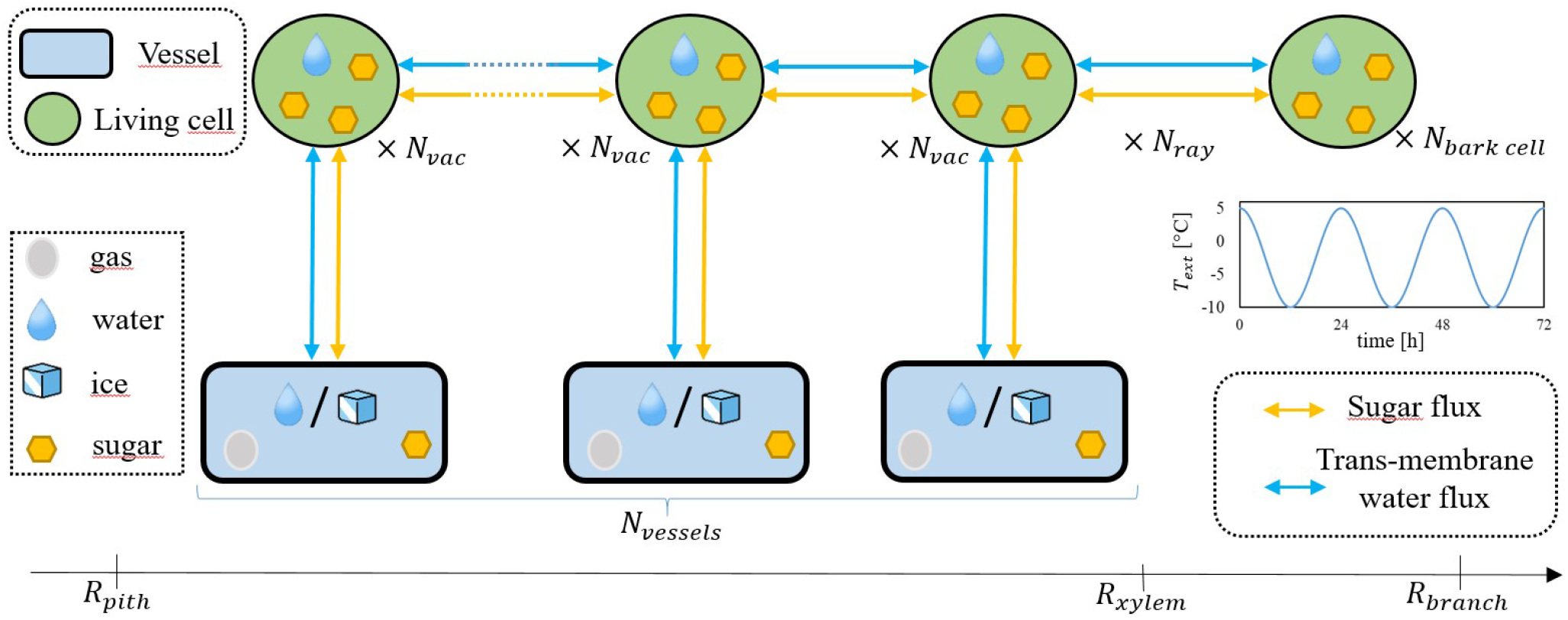
Structure of the model. Depending on the type of element and its enthalpy level, water is assumed to be in different phases (solid, liquid, gas). Water fluxes (blue arrows) and sugar fluxes (yellow arrows) occur between different anatomical elements across cell membranes. Scaling coefficients (*N*_*vac*_, *N*_*ray*_, *N*_*bark cell*_, *N*_*vessels*_) are used to obtain a more accurate anatomical description.

Elastic living cells, i.e., bark cells and VACs, are assumed to contain only water and soluble sugar. We therefore assume that intracellular ice does not form in the temperature range we study. Rigid vessels contain sugar, liquid water or ice depending on local temperature, and gas. This gas compresses or expands in response to water flows entering or leaving the vessels, thus creating pressure variations according to the ideal gas law, as done in Ceseri and Stockie (2013); Graf et al. (2015).

Heat transfer and phase changes are calculated at the tissue scale and driven by external temperature variations. Vessel sugar content impacts tissue-scale phase change through freezing point depression (FPD).

Water fluxes occur between the different elements (blue arrows in figure 1). These water fluxes are driven by the differences in water potential (hydrostatic/turgor, osmotic, cryostatic) across cell membranes. For each elastic compartment (VACs, bark cells), the balance of water fluxes results in volume changes, which are then used to calculate changes in tissue dimensions, as well as changes in turgor and osmotic potential.

The only difference with our previous work is that sugar quantities are now assumed to vary with time. Sugar fluxes occur between the different elements (yellow arrows in figure 1). For simplicity, these fluxes are assumed to come from passive diffusion, i.e., they are proportional to the concentration gradient between two successive elements. These variations in sugar quantities are then used to calculate sugar concentrations in living cells and vessels, which impacts osmotic potential, thus generating water fluxes, and vessel FPD.

### Mathematical description of the numerical model

#### Anatomy

The anatomical description used in the model is shown in figure 1. Vessels are arranged regularly along the ray, with the vessel number, *N*_*vessels*_, computed using a linear vessel density, *lvd* and the size of the xylem tissue:

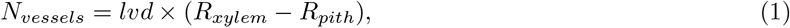

where *R*_*xylem*_ and *R*_*pith*_ are the xylem and the pith radius, respectively. Each vessel has a given number of VACs associated with it, *N*_*vac*_, calculated using a ratio of the vessel-VAC exchange area to the projected VAC area:

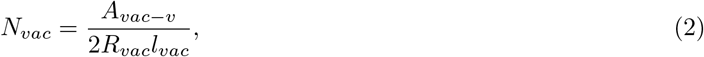

where *A*_*vac*−*v*_, *R*_*vac*_ and *l*_*vac*_ are the vessel-VAC exchange area, the VAC radius and VAC length, respectively. The number of parenchyma rays is computed using a tangential ray density, *trd*, and the branch diameter, *R*_*branch*_:

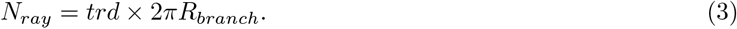

The number of bark cells connected to the parenchyma rays is

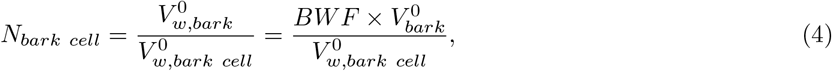

with 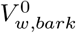 the initial volume of water in the bark accessible from the rays, equal to 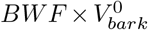, with *BWF* the bark water fraction accessible from the rays, and 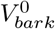 the initial bark volume. 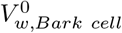 is the initial bark cell water volume.

#### Heat transfer and phase change

Heat transfer and phase change are calculated at the tissue scale through the heat equation in a 1D axi-symmetric model in cylindrical coordinates:

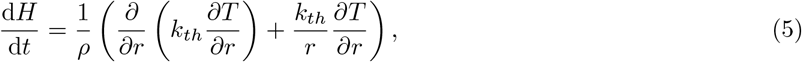

where H is the enthalpy, T the temperature, *k*_*th*_ the thermal conductivity and *ρ* the density. This equation is used for *r*, the radial coordinate, in]*R*_*pith*_; *R*_*xylem*_[and completed with the following boundary conditions (Graf et al., 2015):

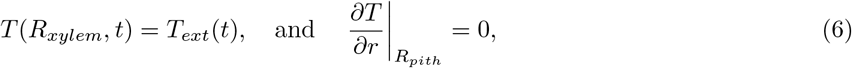

where *T*_*ext*_(*t*) is the external air temperature. Equation (5) must be completed by thermodynamic relationship between *H* and *T* as well as between *H* and the physical properties (density and thermal conductivity) in order to account for phase change. The procedure we used is explained in Bozonnet et al. (2023). The phase change temperature at the tissue scale is computed locally based on 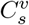, the local vessel sugar content:

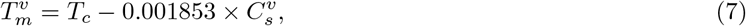

with *T*_*c*_ = 273.15K. Note that Eq. (7) is only valid for water. More details on the implementation can be found in Bozonnet et al. (2023).

#### Water fluxes

Water fluxes between elements are computed using Darcy’s law. Along the parenchyma ray, the fow rate d*V*_*ray*_*/*d*t* is:

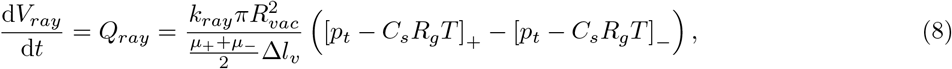

where *k*_*ray*_ is the ray water permeability, Δ*l*_*v*_ is the distance along the ray between two vessels, *µ* is the dynamic viscosity of the water and sugar solution computed locally using the law given in Chenlo et al. (2002), *p*_*t*_, *C*_*s*_ and *T* are the living cell (VAC or bark cell) turgor pressure, sugar concentration and temperature, respectively. The + and − signs represent a differentiation along the ray from the inside to the oustide of the stem (up to the bark cells). *Q*_*ray*_ is positive for water fluxes going towards the inside of the branch. Between one vessel and one corresponding VAC, the flow rate d*V*_*vac*−*v*_*/*d*t* is

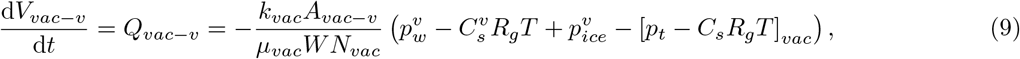

where *k*_*vac*_ is the vessel-VAC membrane water permeability, *W* the vessel-VAC wall thickness, and *p*^*v*^ the vessel water pressure. *Q*_*vac*−*v*_ is positive for water fluxes going towards the vessels. The cryo-suction pressure induced by vessel freezing is computed at each vessel location as (Loch, 1978; Beck et al., 1984)

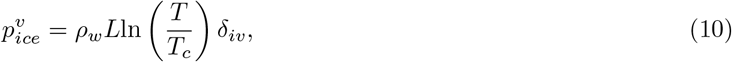

where *ρ*_*w*_, *L*, and *δ*_*iv*_ are the water density, water latent heat of fusion, and vessel ice volume fraction, respectively. Eq. (9) implies that cryo-suction will draw water in a vessel from its VACs once this vessel is frozen.

#### Pressure-volume relationships in living cells

In living cells, the balance of water fluxes results in volume changes. For the VACs, the changes in water volume, d*V*_*vac*−*v*_*/*d*t*, is

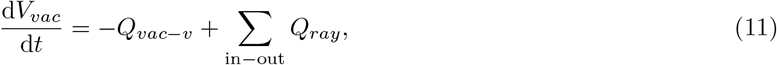

where the second term on the right hand side represents the balance of fluxes entering/leaving the VAC from/to the ray. Between the xylem tissue and the bark, the total water flux is

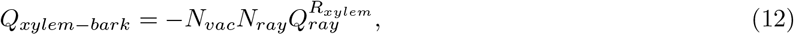

where 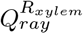 is the water flux computed between one bark cell and the VAC closest to the bark. This water flux is rescaled by the number of bark cells to obtain the volume change at the bark cell scale, *V*_*barkcell*_ :

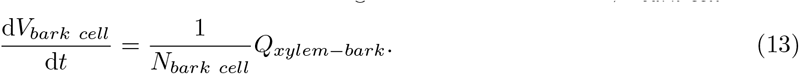

These living cells’ volume changes are related to turgor pressure variation through (Steudle et al., 1977)

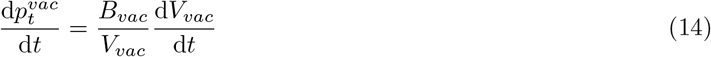

for the VACs, and

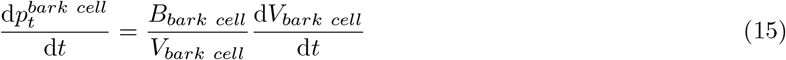

for the bark cell. In the previous two equations, *B*_*vac*_ and *B*_*bark cell*_ are the VAC and bark cell elastic modulus, respectively. We use the procedure introduced in Bozonnet et al. (2023) to account for turgor loss. Volume changes also result in osmotic pressure changes through changes in sugar concentration, which are also related to changes in sugar content:

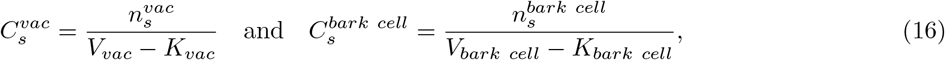

where 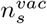 and 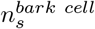 are the variable sugar quantities in the VAC and bark cell, and *K*_*vac*_, *K*_*bark cell*_, are the cell volume where no sugar can be contained (certain cell organelles), respectively. These changes in sugar concentration also result in living cell FPD for the VACs and the bark cells, with an equation similar to Eq. (7).

#### Pressure-volume relationships in vessels

Vessels contain gas that compresses or expands depending on water fluxes leaving or entering vessels. Following Ceseri and Stockie (2013); Graf et al. (2015); Bozonnet et al. (2023), we assume that this gas is contained in one cylindrical bubble located at the center of each vessel. Applying flow rate conservation between the gas/water (or gas/ice) interface and the vessel/VAC membrane, the bubble radius, 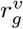, varies as

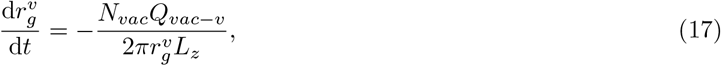

where *L*_*z*_ is a vertical dimension that is introduced for unit consistency but that has no influence on model results. Changes in 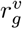 induce changes in gas pressure through the ideal gas law:

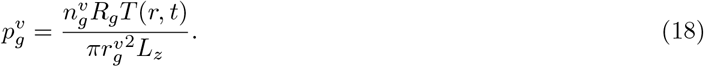

In previous equation, 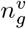 is the gas quantity inside the gas bubble and *R*_*g*_ the ideal gas constant. Finally, the pressure in the liquid water/ice phase is obtained using Laplace equation:

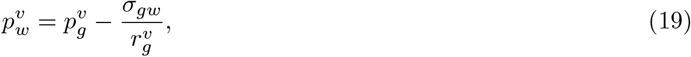

where *σ*_*gw*_ is the liquid water/gas interface surface tension. In the results section, we will also use the evolution of the mean vesssel pressure over all vessels, defined as

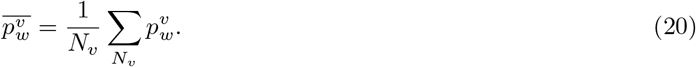

Similarly to living cells, water volume and sugar content changes in vessels also result in sugar concentration changes:

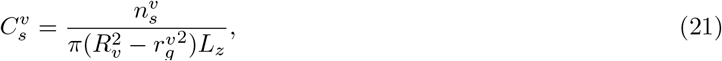

where 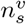 and *R*_*v*_ are the variable vessel sugar quantity, and vessel radius, respectively. These changes in vessel sugar concentration impact phase change at the tissue scale through Eq. (7). We will also use in the results section the average vessel sugar concentration, 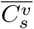, defined in a way similar to Eq. 20.

#### Sugar fluxes

Sugar fluxes occur due to passive diffusion between elements. Each vessel sugar content is computed as

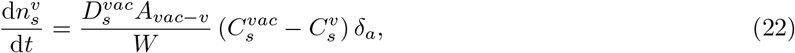

with 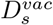 the sugar diffusion coefficient between one vessel and one VAC, and *δ*_*a*_ is an activation coefficient. *δ*_*a*_ goes linearly from 0 at -0.5°C to 1 at 0°C, and outside this interval it is equal to 0 for lower temperature and 1 for higher temperature, hence progressively blocking sugar diffusion at negative temperature. It appeared essential for numerical stability to block sugar diffusion at negative temperature, as the code had difficulties to converge at negative temperature when ice was blocked in vessels whereas sugar fluxes would still induce water flows in-between living cells. Each VAC sugar content is computed as

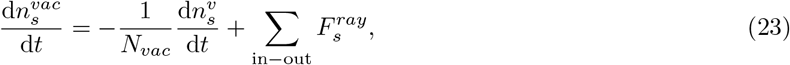

where the first term on the right-hand side represents the sugar flux leaving the VAC towards its vessel, and the second term represents the sum of sugar fluxes leaving or entering each VAC to/from the ray. These fluxes are computed as

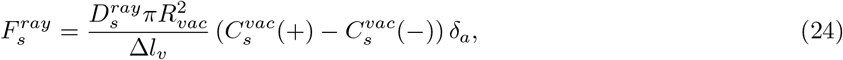

where 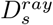 is the sugar diffusion coefficient across the ray, and the + and − signs represent a differentiation along the ray from the inside to the oustide of the stem (up to the bark cells). 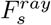 is positive for sugar transport towards the inside of the stem. The bark sugar content is computed as

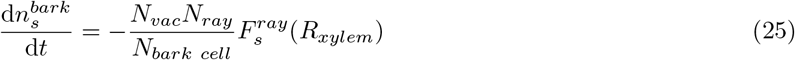

#### Diameter changes

Finally, diameter changes are obtained from living cell volume changes, Eqs. (11) and (13). The total volume of water in VACs is computed at each instant:

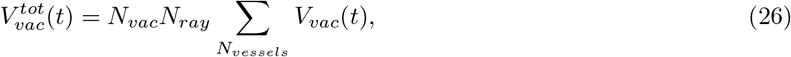

with the volume variation equals to

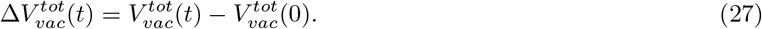

The xylem diameter, considered as a cylinder, is computed as

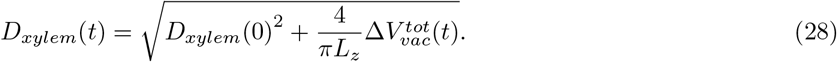

At each instant, the volume of the bark tissue is equal to the initial volume minus the volume lost by dehydration:

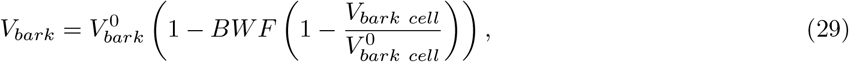

from which the stem diameter, considered as a cylinder, can be computed:

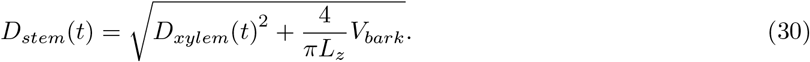

#### Numerical resolution and parameter choices

The model is implemented in the Matlab software version R2018a (MATLAB, 2018). Spatial discretisation of Eq. (5) is ensured using the finite difference method. The system of differential equations formed by equations (5), (8), (9), (11), (13), (14), (15), (17), (22), (23) and (25) is advanced in time using Matlab’s variable order *ode*15*s* solver based on numerical differentiation formulas, which is specifically designed for stiff equations, with a maximal time step of 1*s*, which is a sufficient value to resolve any stiffness in the problem under study. The other equations are state equations computed at each time step. Note that we verified the implementation of Eq. (5) using an analytical solution (Prapainop and Maneeratana, 2004) for a 1D freezing-front propagation (Stefan problem, R^2^ *>* 0.9999), and using a reference finite element solver (Comsol Multiphysics (COMSOL, 2020)) for the 1D axi-symmetric implementation (R^2^ = 0.9998). The reference result presented in figure 2 takes a computational time of around 10 minutes on a Dell Latitude 7490 with 1.7 GHz quad-core Intel i5 processor. The source code can be downloaded at https://github.com/cyrilbz/pressurebuildup.

**Figure 2:**
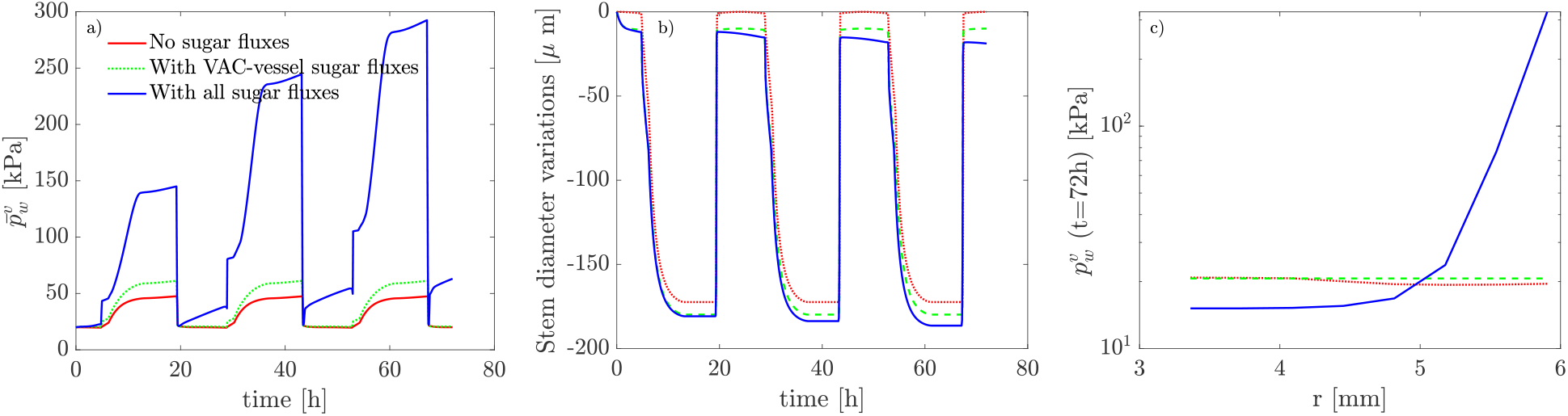
Effect of sugar fluxes on mean pressure (a), stem diameter changes (b), and pressure field at the end of the simulation (c). In (c), *r* is the radial coordinate. Continuous red line: 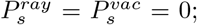 green dashed line: 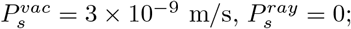 continuous blue line: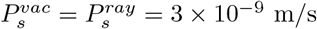.

All model and state variables are regrouped in table 1. All parameters are in table 2. All values are either justified based on the literature, have been specifically measured, or calibrated and justified in our previous work (Bozonnet et al., 2023), except the initial sugar content in living cells, and the sugar diffusion coefficients. The initial sugar content in living cells has been estimated based on measurements on whole stems in Charrier et al. (2013). The starting values for the diffusion coefficients have been computed using a solute permeability coefficient 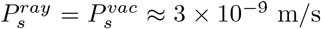 (Gunning, 1977; Tyree et al., 1994), which leads to 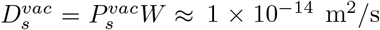 and 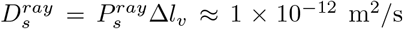. We note that the permeability coefficients in the reference we used correspond to solute flow across roots, hence the actual values might be underestimated.

**Table 1:**
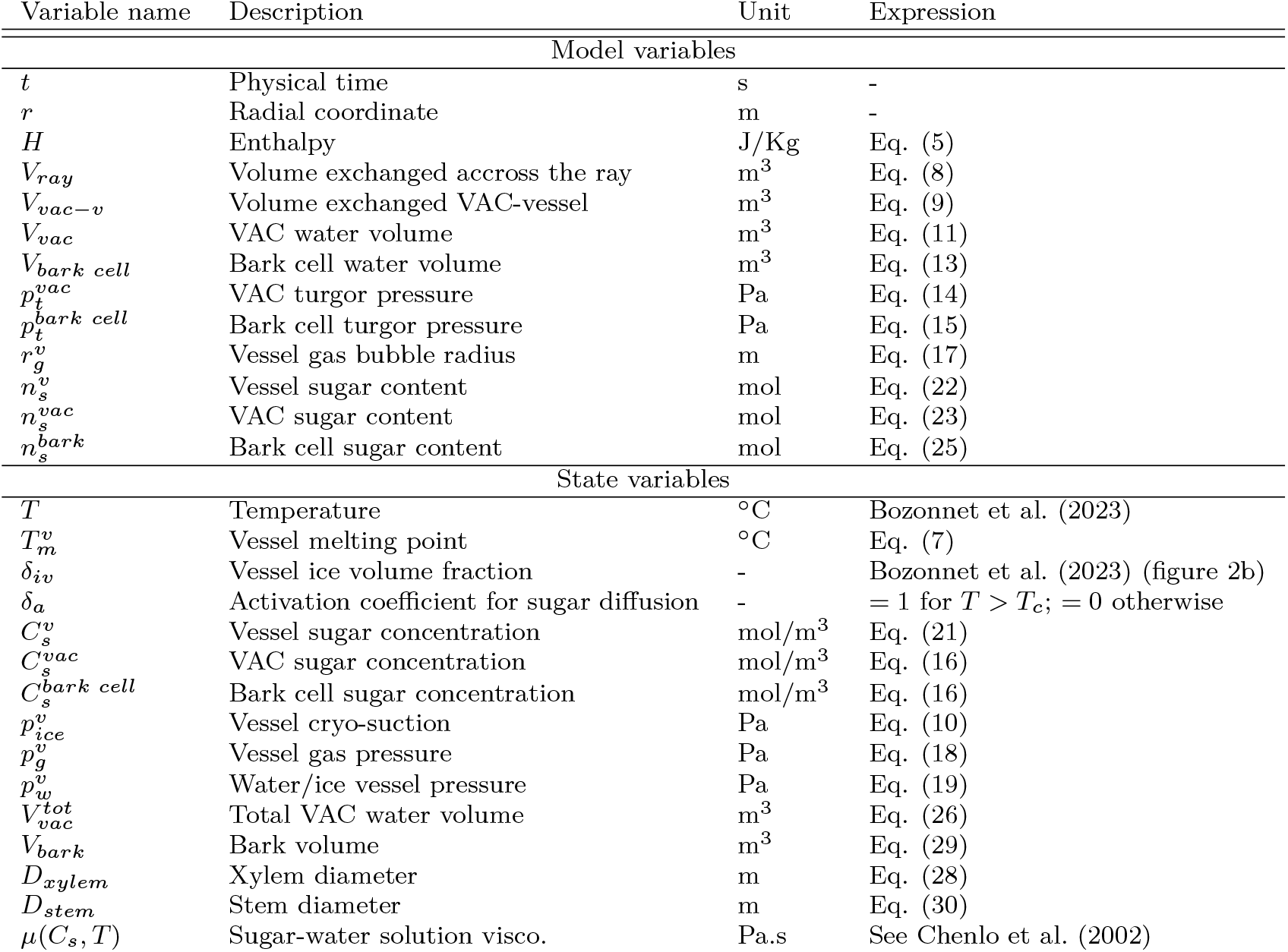
Model and state variables.

**Table 2:**
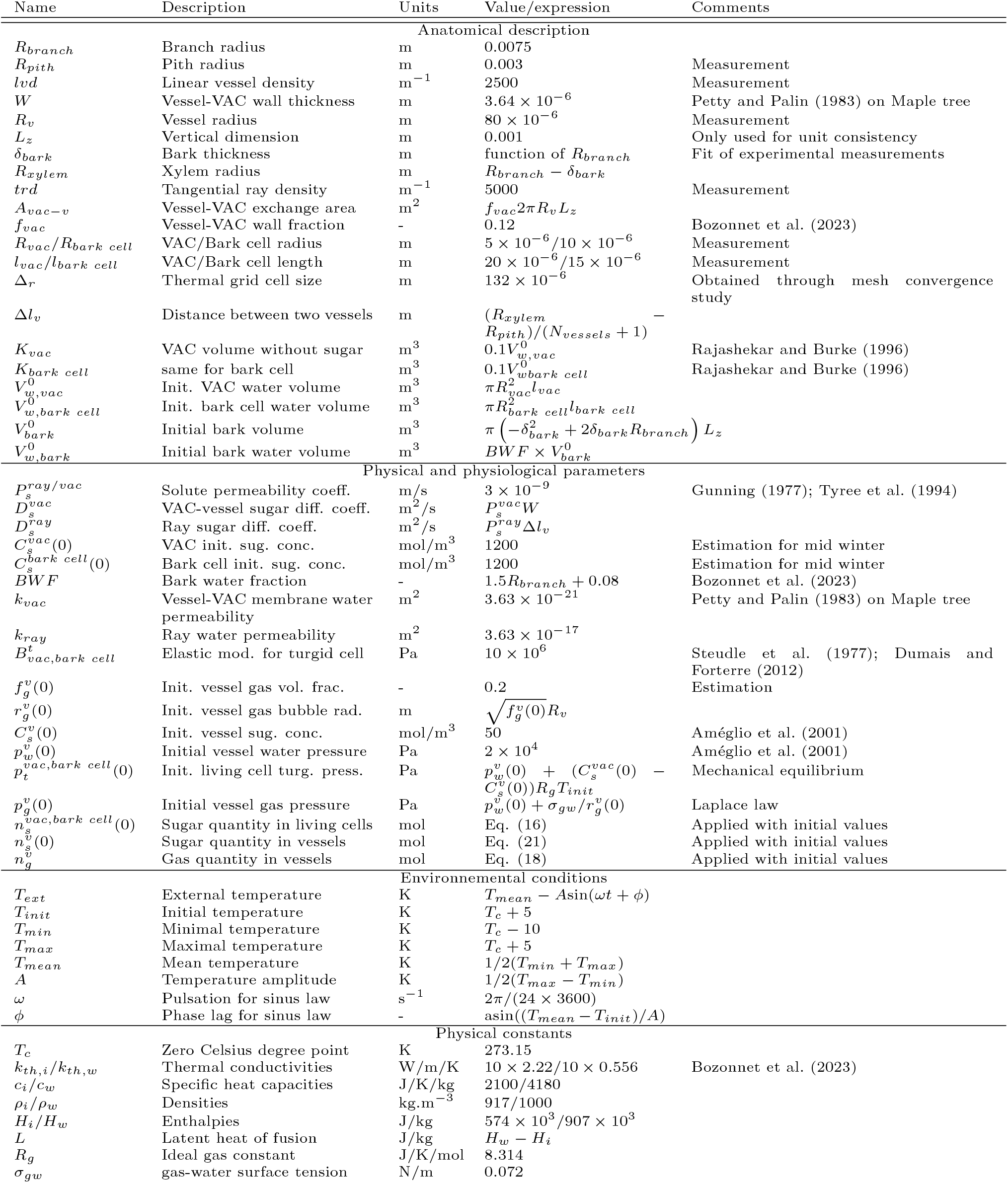
Parameter list, description & values. Measurements and estimations were done for *Juglans regia* stems.

For simplicity, we further assume that all living cells have the same mechanical properties and the same initial sugar concentrations. However, the model is already capable of handling different parameter values between VACs and bark cells.

## Results

In this section we present the results obtained using the model described previously. Unless stated otherwise, all parameter values are presented in table 2. In the following, and unless stated otherwise, the term pressure always refers to the vessel pressure in the liquid water/ice phase, 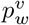 .

### Effect of sugar fluxes

In this section we describe the model results obtained with or without sugar fluxes. The stem undergoes a three-day period of continuous temperature fluctuations, ranging between +5°C and −10°C, occurring in 24-hours cycles, as shown in figure 1. The expression for *T*_*ext*_(*t*) is given in table 2.

Without sugar fluxes (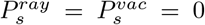; continuous red line), the mean pressure (figure 2a) shows an alternating sequence of increases at freezing and drops at thawing. As explained in Bozonnet et al. (2023), this is due to water going from living cells to xylem vessels under the influence of the low ice potential in vessels and reversing at thawing. The effect of these water flows can be seen in stem diameter changes (figure 2b): at freezing we observe a shrinkage in diameter, followed by a swelling at thawing. Pressure and stem diameter changes are fully reversible. In the end of the simulation the pressure distribution (figure 2c) is nearly homogeneous across the stem, i.e., pressure values are equal for all radial positions, and the mean pressure is equal to its initial value. When sugar fluxes between VAC and vessels are included (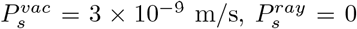; green dashed line), we observe a slight increase of the final pressure compared to the case where these fluxes were missing (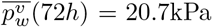 compared to 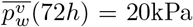, see figure 2a). The maximun pressure in the frozen state increases, but more significantly (+15kPa). Stem diameter changes are also affected: compared to the previous case, the maximun shrinkage at freezing increases, i.e., the minimal diameter decreases, and at thawing the diameter does not come back to its initial value (figure 2b). The radial pressure profile (figure 2c) does not show much difference with the previous case. Note that increasing the number of VACs per vessel (*N*_*vac*_ = 2260 compared to *N*_*vac*_ = 300 in the present case) presents only a slight vessel pressure increase after 72 hours (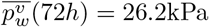, results not shown).

When radial sugar fluxes are added (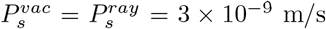; continuous blue line), the mean vessel pressure shows a much larger increase at the end of the simulation: 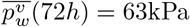 (figure 2a). It progressively increases in both states: in the frozen state, the maximum pressure increases all over the simulation, as well as in the thawed state, where sugar fluxes are active. Stem diameter variations (figure 2b) also show differences in both states: in the frozen state, the maximum shrinkage progressively increases over the three cycles, and in the thawed state, the maximum diameter decreases with time. The radial vessel pressure profile shows spectacular differences with the other two previously cases (figure 2c): vessel pressure is slightly lower (−25%) than previously near the pith (lowest *r* values), and significantly higher (up to +1530%) near the bark (highest *r* values).

### Effect of sugar permeabilities

In figure 3a we show the effect of both sugar permeability coefficients (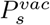 and 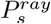) on the pressure level after 72*h*. We consider three cases: one case where 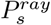 is kept equal to its original value (3 *×* 10^−9^ m/s) and only 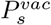 varies (continuous line), and two cases where 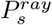 is kept constant and only 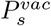 varies, for 2 values of *P*^*vac*^ (dashed lines).

**Figure 3:**
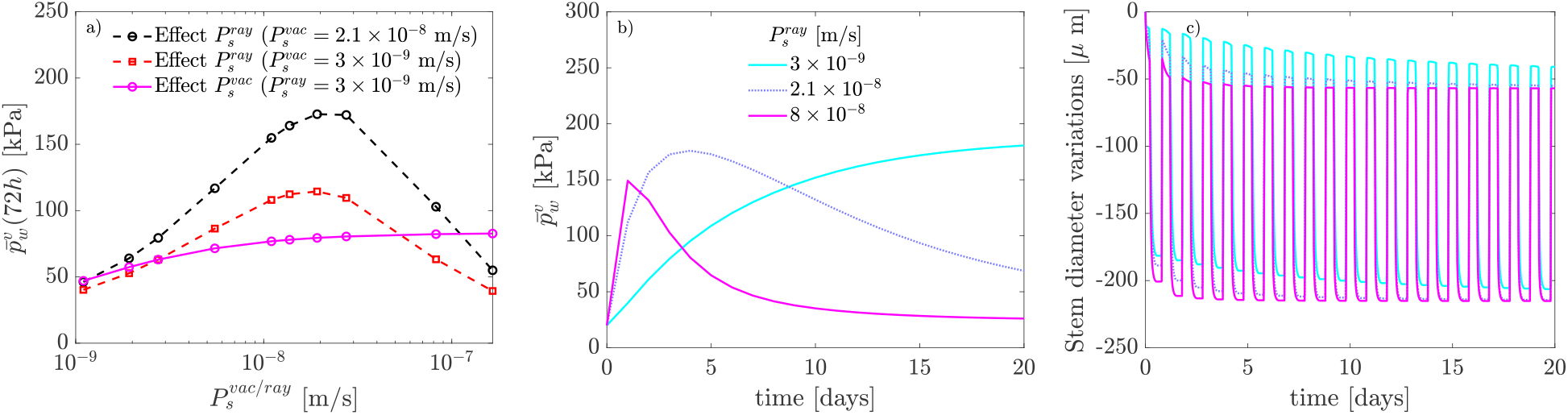
Effect of sugar permeabilities (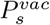 and 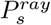) on the mean vessel pressure after 72 hours (a). Effect of the ray sugar permeability 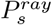 on the mean pressure long term evolution (b), and on stem diameter changes (c). Note that in b only one data point every 24 hours (in thawed state) is shown to enhance readability. In b and c: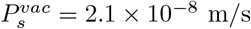. For the legend in figure c, please refer to figure b.

We observe that when 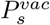 only is increased, the pressure build-up increases too and reaches a plateau at high 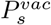. When 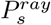 only is varied the pressure build-up is low for extreme 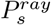 values and passes through a maximum value. This is valid for both data series at varying 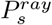, with the data serie for the highest 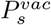 showing the highest pressure levels.

In figure 3b we show the effect of the ray sugar permeability on the long term evolution (20 days) of the mean pressure. The case with 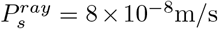 shows the fastest pressure increase followed by a decline after only 2 days. After 10 days, the case with 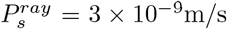, the lowest 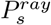 value in this figure, generates the highest pressure level. For a sufficiently long simulation, and for the parameter range considered here, the maximal mean pressure level reached during a simulation decreases with an increase in ray sugar permeability. In figure 3c we show the effect of the ray sugar permeability on the long term (20 days) variations of stem diameter. One can see that increasing the permeability decreases the diameter, both in frozen and thawed states.

### Spatio-temporal vessel pressure variations

In figure 4a we draw from figure 3b the changes in mean vessel pressure for the case with 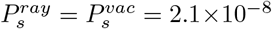, with a custom color code that represents the time course. We chose this case as it shows a pressure build-up followed by a decline. In figure 4b we show for the same case the vessel pressure profile as a function of the radial coordinate and for different instants of the simulation (one profile every 24 hours) with the same color code. Similarly to figure 2c and compared to the initial value, the pressure decreases near the pith and increases near the bark. One can see that in only 2 days the vessel pressure reaches its maximal value near the bark. Then the pressure curve progressively spreads towards the interior of the stem. For a much longer simulation, the vessel pressure will become homogeneous along the radius (results not shown), similarly to cases with no ray sugar fluxes in figure 1a, but with a slightly higher pressure value.

**Figure 4:**
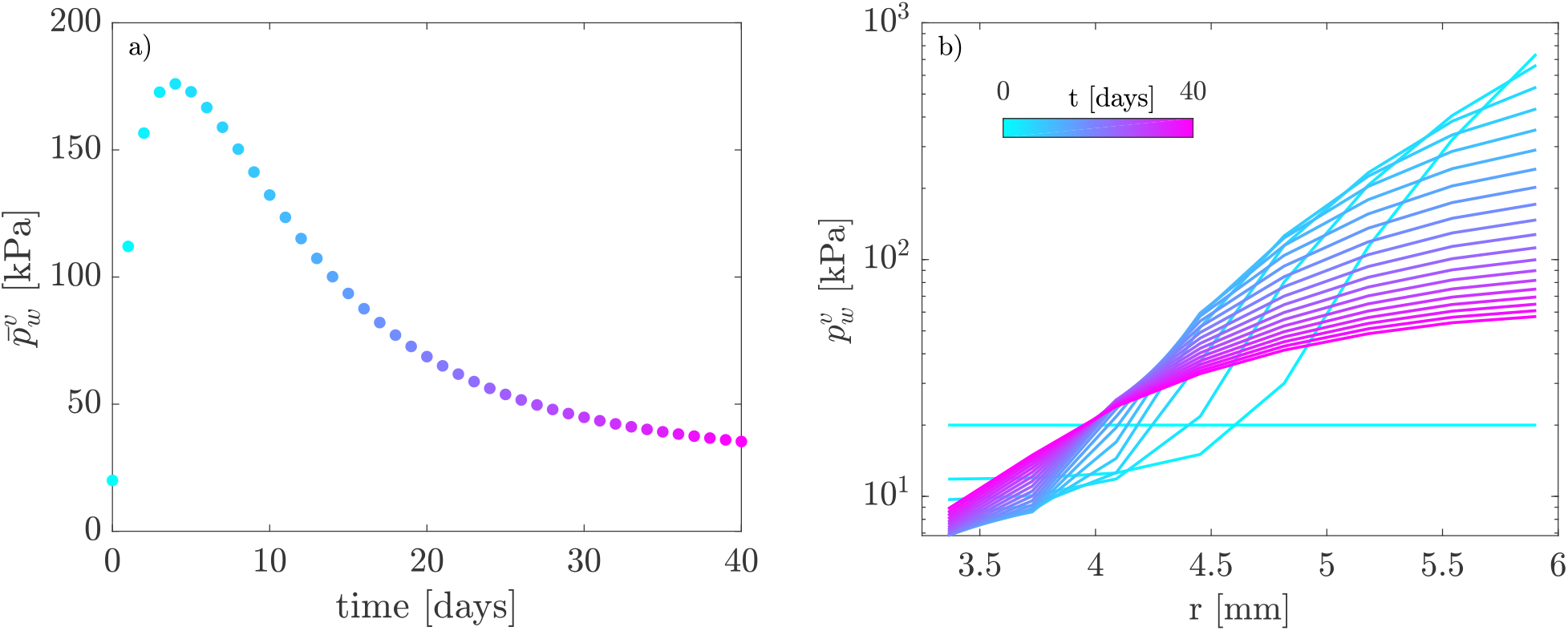
Spatio-temporal variations of vessel pressure for case 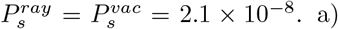 Mean pressure signal (reproduced from figure 3 with a custom color code), for color interpretation see the scale in figure b. In a, only one data point every 24h is shown to enhance readability. b) Radial pressure profile every 48h.

### Effect of initial sugar content and validation

Figure 5a illustrates the effect of the initial sugar concentration in living cells on the mean pressure for two sugar permeability values. The initial sugar concentrations were chosen to reflect the potential changes in soluble carbohydrates content across the winter season (Charrier et al., 2018), and with the treatments applied to the trees (defoliation, exposure to low or high temperature). We remind the reader that all living cells initially have the same sugar content. We observe an increase of the mean pressure with the initial sugar content. This increase is even higher at higher sugar permeability. For low initial sugar concentration (200mol/m^3^) the difference between both cases is only 8kPa, whereas it reaches 160kPa for the highest initial sugar concentration (1600mol/m^3^).

**Figure 5:**
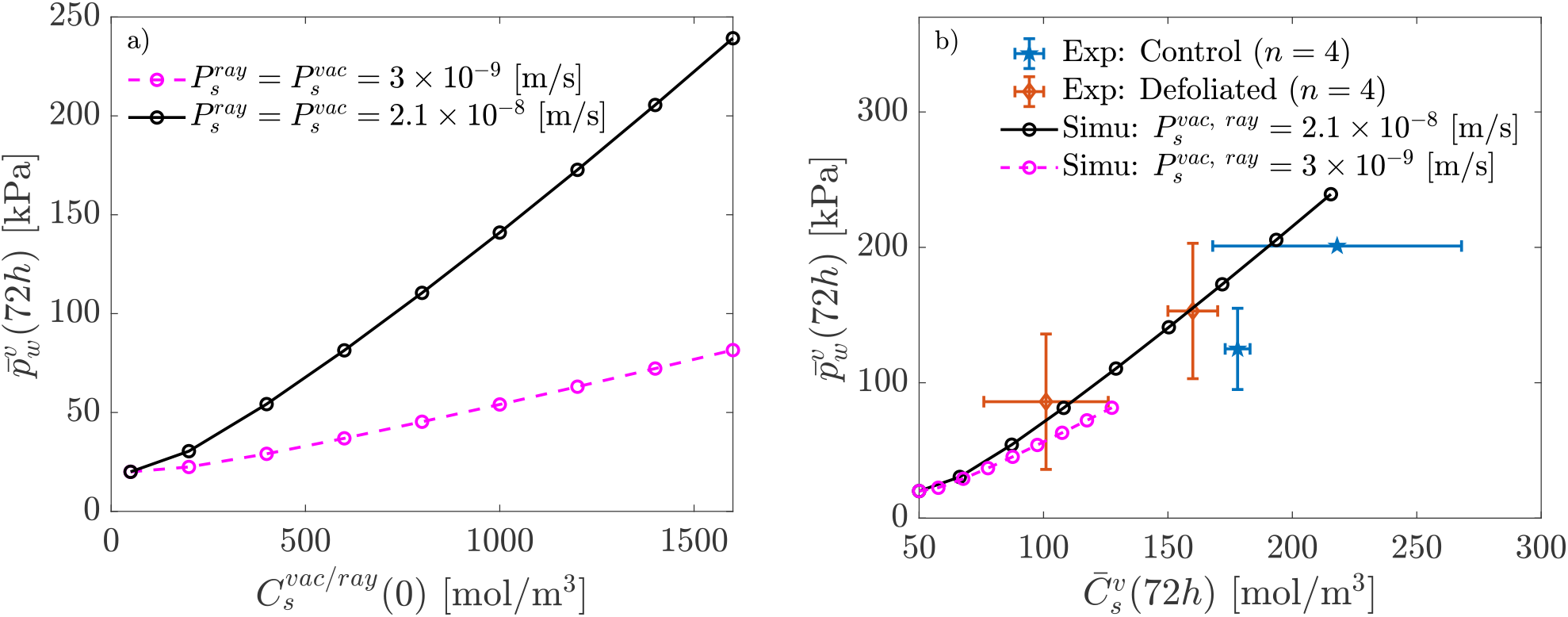
a) Effect of initial living cell sugar concentration on the mean vessel pressure after 72h. b) Link between mean vessel pressure and mean vessel sugar content after 72h + comparison with experiments from Améglio et al 2001 (after 72h too).

In figure 5b we show the relation between the mean vessel sugar concentration and the mean vessel pressure, both after 72h, for the cases presented in figure 5a. In the same figure we also show the experimental results from Améglio et al. (2001), for stems that were defoliated during summer and the control group. The mean pressure level rises with the increase of the mean vessel sugar concentration for both the model and the experimental data. For the greatest permeability coefficient, both the average sugar concentration and mean pressure reach significantly higher values. Model results and experiments have similar order of magnitude for both quantities.

## Discussion

### Mechanism behind pressure build-up

As we shown in figure 2a, in our model sugar fluxes across the parenchyma ray are absolutely essential for pressure build-up to occur. They also dramatically change the pressure radial distribution within the stem, with the pressure decreasing near the pith and increasing elsewhere (figure 2c). They are initially induced by the sugar fluxes from VACs to vessel, that decrease the VACs sugar concentration. These ray sugar fluxes act on the pressure through two mechanisms. First, the sugar flux from the bark cells to the vessels (through the VACs) induce a flow of water from the bark cells towards the vessels, thus increasing the pressure. Secondly, because of the spatial distribution of the vessels along the parenchyma rays, the vessels close to the bark are preferentially loaded with sugars. This creates a radial gradient in sugar concentration that induces a water flux from the near pith vessels to the near bark vessels, thus creating the distribution in pressure observed in figure 2c.

The first mechanism, the sugar and water fluxes from the living cells towards the vessels, can also be evidenced through stem diameter changes: a reduction in diameter occurs both in frozen and thawed states. Particularly, the increase in the freeze-induced stem shrinkage is due to the decrease in living cell sugar content, as already shown in Bozonnet et al. (2023). The second mechanism, although not directly observable on stem diameter has a much greater impact on vessel pressure: the case with the highest stem pressure does not have the smallest diameter in the frozen state (see figure 3b and c). We have thus demonstrated that pressure build-up can be due to a transfer of water between vessels, across the parenchyma rays, induced by a radial imbalance in vessel sugar concentration.

For low and high ray sugar permeabilities 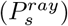, the radial imbalance is greatly reduced, and no pressure rise occurs, except the one due to the water flows coming from living cells, as shown in figure 3a with the effect of 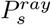. Sugar transportation along the ray alone is not sufficient: if VAC to vessel transport is too low, no pressure accumulates, as observed in 3a, with the effect of 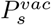. Note that, in the short term, there will necessarily be no sugar radial imbalance, thus no pressure rise. This is also true for sufficiently long simulations for which the sugar concentrations will become homogeneous across the stem (figure 3b). The duration required to achieve this homogenisation, as well as the duration required to reach the maximum pressure build-up, is contingent on the ray sugar permeability values.

The spatio-temporal variations of the vessel pressure give even more insights into the process. Figure 4b enlightens that at short times there is a pressure decrease near the pith and an increase near the bark, indicating a flow of water in-between vessels through the parenchyma rays. The decrease near the pith continues while the pressure profile in the vicinity of the bark progressively flattens. The mean pressure reaches its maximum during this part of the process (figure 4a). Eventually the pressure near the pith progressively starts to increase while the pressure profile tends towards homogeneity. For sufficiently long simulations, pressure values will be equal for all vessels, with a value slightly higher than for the case where no ray sugar fluxes were included in the processes (figure 2).

### Comparison with experiments

Stems that underwent early defoliation showed low pressure build-up compared to controls (Améglio et al., 2001). Early defoliation can indeed reduce the amount of stored carbohydrates in living cells, thus preventing them to hydrolize starch during winter in order to increase their sugar concentration (Charrier et al., 2018). Similarly, a treatment at high temperature before the experiments was also likely to reduce the accumulation of soluble sugar in living cells. Both of these treatments lowered the measured pressure level in Améglio et al. (2001). The model predicts the same relation between the initial sugar concentration and the mean vessel pressure level (figure 5a): a decrease in the concentration leads to a decrease in pressure. The effect of the initial sugar concentration is even greater at high sugar permeability, which is fully expected as higher permeabilities lead to higher presure (figure 3a).

Finally, we have validated the model by comparing two of its outputs, the mean vessel sugar concentration and the mean vessel pressure after 72h, against the measured sap osmolarity and xylem pressure after 72h of experiments. We emphasis that both of these quantities are results, either from the model or the experiments, and not inputs or controlled parameters. The comparison is thus extremely favorable, as similar orders of magnitude are reached between the model and the experiments for both of these quantities, especially for the model results at high sugar permeability coefficients. This shows that the initial permeability coefficients we used were probably underestimated.

We note that the external temperature between the simulations and the experiments were not exactly the same: in Améglio et al. (2001), stems that underwent a high temperature treatment before the experiments were exposed during the freeze-thaw cycles to a maximal temperature up to 18°C, while the other stems were exposed to a maximal temperature of 1.5°C. In the simulations, we chose a maximal temperature of 5°C, that we estimate as being a threshold above which H^+^/sugar co-transport will bring sugar from the vessels back to the living cells and in-between living cells (at counter gradient). This mode of transport is not included in the model, but including it would be a way to go further in the exploration of temperature effects on stem pressure build-up. This would also be a way to verify if the current understanding of these temperature effects, as being the result of a balance between the H^+^/sugar co-transport and diffusion, is correct.

Similarly, starch-soluble sugar inter-conversion is not included in the model as we expect it to occur on a timescale longer than 3 days (Charrier et al., 2018). It can however have an impact on longer time scales. Including it in the model would be a way to dynamically compute the initial living cell sugar concentration, as a function of the environmental conditions and the tree’s carbohydrate reserves. This way, the effect of the different experimental treatments (defoliation, low/high temperature exposure) could be simulated directly within the model and not modelled with a varying initial sugar concentration.

In the model from Graf et al. (2015), pressure build-up, as we defined it in the introduction, occurs due to irreversible root absorption during freezing or thawing events in the stem. This is different from our model results and the experiments of Améglio et al. (2001), where it occurs during the day (at slightly positive temperature, after thawing and before freezing), and, particularly, in the absence of any connection with the root system. In Graf et al. (2015), freeze-thaw cycles are essentials for this build-up to occur, as in the experiments of Améglio et al. (2001). Our model does not reproduce such synergetic effect of freeze-thaw cycles: a case without freezing temperature shows the same pressure build-up as a complete case. Both models predict an increase in vessel pressure with sugar content, although in Graf et al. (2015) the sugar content does not change with time. Only our model reaches orders of magnitude that are consistent with the experimental results for both the final vessel pressure and vessel sugar concentration.

Although water fluxes between crown and roots are occurring in field experiments (Charrier et al., 2017), and are essentials for maple sap harvest (Tyree, 1984), we do not think that adding them in our model is the path to follow to obtain this synergetic effect of freeze-thaw cycle. This would indeed requires a connection with the rest of the tree, whereas following Améglio et al. (2001) it is not needed for pressure build-up to occur. It is possible that our omission of sugar transport at negative temperatures and our assumption of a constant quantity of gas in vessels, 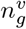, contributed to this lack of synergetic effect. It is well-established that freeze-thaw cycles create gas bubbles that may fill an entire section of the vessel, see references in the introduction. One could hypothesize that repeated freeze-thaw cycles could raise the value of 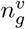, thereby affecting the dynamics of pressure build-up. One other missing ingredient could be the starch to sugar conversion in living cells, which could still occur at slightly negative temperature, but as said previously we do not expect it to have an impact on such a short timescale. It would be worth investigating further this point to understand better the effect of freeze-thaw cycles on pressure build-up and how it could be impacted by a changing climate.

The pressure build-up mechanism we have highlighted leads to a pressure drop near the pith and a pressure rise elsewhere. In the context of embolism recovery, this means that this mechanism cannot repair embolism in the vicinity of the pith. We currently have no experimental data to validate such consequence. Such data could be obtained by scanning (X-ray microtomography) walnut branches before and after stem pressure generation to precisely locate the places where embolism recovery occurs, such as done for example on birch and maple tree in Robinson et al. (2023). This heterogeneous repair might however be related to the transition from sapwood to heartwood due to tyloses formation in embolized vessels (Kozlowski and Pallardy, 1997; Barnett, 2004). Another way to validate the mechanism would be to repeat the experiments of Améglio et al. (2001) with stem samples that have their bark removed and intact ones. The absence of pressure build-up for stems with their bark removed would be a strong argument to validate the mechanism.

Compared to our previous model that did not generate any pressure build-up (Bozonnet et al., 2023), the only additional mechanism in the present one is the transport of sugar by diffusion in-between living cells and between living cells and vessels. The reason why some species are (or not) able to increase the pressure in their branches might therefore lie in this ability to transport sugar during winter. In the walnut tree, this is regulated by specific proteins in cell membranes, that could be less abundant during winter in other species. This might also depend on anatomical features such as the number of vessel associated cells, or the parenchyma rays anatomy.

In addition, it is worth investigating in the future whether this mechanism is relevant to the development of winter stem pressure in maple trees and the harvesting of maple sap. Particularly, one could start by coupling our work with recent modelling efforts on maple tree (Graf et al., 2015; Zarrinderakht et al., 2024). One major difference between maple and walnut trees lies in the need, when modelling pressure changes in maple tree, to include a hydraulic connection as well as an osmotic barrier between vessels and fibers. This is indeed required to reproduce the pressure drop observed at freezing inception in maple trees (Milburn and O’Malley, 1984; Cirelli et al., 2008; Ceseri and Stockie, 2013; Graf et al., 2015; Zarrinderakht et al., 2024), whereas the opposite is observed in walnut trees and all other species (Robson and Petty, 1987; Améglio et al., 2001). This pressure drop at freezing favours water entry from the roots while vessels are still in the liquid state. Water exchange with the roots, even though, in our opinion, not a key ingredient in the pressure build-up as discussed previously, can therefore have a much greater impact in maple tree compared to walnut tree. In both species, however, water fluxes between the crown and the roots could be directly driven by cryostatic suction, which is not taken into account in any of the existing models.

## Conclusion

The initial aim of this work was to develop a mechanistic model capable of simulating the winter pressure build-up in walnut stems. We have shown that the pressure build-up can be explained by a transfer of water between vessels via the parenchyma rays, induced by a radial imbalance in sugar concentration.

Among the various features listed in the introduction, this mechanism succeeds in: generating a pressure build-up for stems disconnected from the rest of the tree, quantifying the relationship between pressure build-up and xylem sap osmolarity, and showing the effect of experimental treatments (defoliation, low/high temperature exposure) through the influence of the initial living cell sugar concentration. Temperature effects on vessel pressure are partially captured due to the lack of H^+^/sugar co-transport in the model. The model does not yet capture the synergetic effect of freeze-thaw cycles on pressure build-up, which needs to be investigated in the future.

Finally, we have outlined two experiments to validate the pressure build-up mechanism we have identified. Both of these experiments could validate two crucial aspects of the mechanism: it leads to a heterogeneous embolism repair, and it requires sugar fluxes coming from the bark.

## Data availability statement

The source code used to generate the data of the present paper can be downloaded at https://github.com/cyrilbz/pressurebuildup. Any result of the present paper can be reproduced using this code.

## Acknowledgments

The ANR, through the ACOUFOLLOW project (ANR-20-CE91-0008), is acknowledged for the funding of this project.

